# Within-species variation in the gut microbiome of medaka (*Oryzias latipes*) is driven by the interaction of light intensity and genetic background

**DOI:** 10.1101/2023.02.17.528956

**Authors:** C. Evangelista, S. Kamenova, B. Diaz Pauli, J. Sandkjenn, L.A. Vøllestad, E. Edeline, P. Trosvik, EJ. de Muinck

## Abstract

Unravelling evolution-by-environment interactions on the gut microbiome is particularly relevant considering the unprecedented level of human-driven disruption of the ecological and evolutionary trajectories of species. Here, we aimed to evaluate whether an evolutionary response to size-selective mortality influences the gut microbiome of medaka (*Oryzias latipes*), how environmental conditions interact with the genetic background of medaka on their microbiota, and the association between microbiome diversity and medaka growth-related traits. To do so, we studied two lineages of medaka with known divergence in foraging efficiency and life history raised under antagonistic size-selective regimes for 10 generations (i.e. the largest or the smallest breeders were removed to mimic fishing-like or natural mortality). In pond mesocosms, the two lineages were subjected to contrasting population density and light intensity (used as proxies of resource availability). We observed significant differences in the gut microbiome composition and richness between the two lines, and this effect was mediated by light intensity. The bacterial richness of fishing-like medaka (small-breeder line) was reduced by 34% under low-light conditions compared to high-light conditions, while it remained unchanged in natural mortality-selected medaka (large-breeder line). However, the observed changes in bacterial richness did not correlate with changes in adult growth rate or body condition. Given the growing evidence about the gut microbiomes importance to host health, more in-depth studies are required to fully understand the role of the microbiome in size-selected organisms and the possible ecosystem-level consequences.

## Introduction

Over the past decade, the expansion of scientific literature investigating the gut microbiome has been largely motivated by increasing evidence of the microbiome’s role in health maintenance. Studies have highlighted the variety of pathways through which the gut microbiome can play this role – from influencing nutrient uptake and metabolism to regulating immune responses and pathogen susceptibility (Hanning and Diaz-Sanchez 2015, Moran et al. 2019). Gut-associated microbiomes can also incur the consequences of various anthropogenic stressors (Sepulveda and Moeller 2020, Teyssier et al. 2018, Trosvik et al. 2018, Sonnenburg and Sonnenburg 2019, Fouladi et al. 2020, Varg et al. 2021). These anthropogenic stressors may lead to imbalances in the gut microbiome (dysbiosis) with potential implications for individual host fitness. A notable example is the effect of increased temperature leading to reduced gut bacterial diversity in the common lizard (*Zootoca vivipara*), also potentially associated with a reduction in survival (Bestion et al. 2017). Understanding how changes in the gut microbiome are related to host health and survival is essential to fully understand the mechanisms behind anthropogenic impact on wild populations.

In teleosts (bony fish), the genetic background (Bolnick et al. 2014a, Smith et al. 2015) and various environmental factors such as diet composition (Bolnick et al. 2014b, Eckert et al. 2020) and the quality of the surrounding water (Talwar et al. 2018) have been linked to changes in the gut microbiome composition. However, as genotypes and environmental conditions are often interacting, disentangling their respective effects on the microbiome is challenging, and have been mainly studied independently (but see Navarrete et al. 2012, Sevellec et al. 2014, Sullam et al. 2015). Currently, we lack assessments of genotype-by-environment effects on fish microbiome (Spor et al. 2011, Talwar et al. 2018, Piazzon et al. 2020) which limit our understanding about the factors driving variation in gut microbiome assemblages in teleosts, a comparatively under-studied taxa in microbiome science (Sullam et al. 2012).

Size-selective harvesting of wild populations is an acute disturbance factor, driving very fast rates of evolutionary change towards smaller size and faster life history (Darimont et al. 2009, Nusslé et al. 2012, Heino et al. 2015, Sanderson et al. 2022). Simultaneously, harvesting reduces population density and hence also increases resource availability. The evolutionary change driven by size-selective harvesting (van Wijk et al. 2012, Uusi-Heikkilä et al. 2017) has, in turn, the potential to reshuffle trophic interactions within food webs, with individuals from heavily harvested populations tending to display narrower diets (Hočevar and Kuparinen 2021). Assuming that bacterial transmission through diet influences gut microbial diversity, individuals with narrower diets may have less diverse bacterial communities (Sevellec et al. 2014). In fish exposed to fishing, the diet and gut microbiome of fish could be altered by the evolutionary change in size due to size-selective harvesting, the increase prey abundance due to reduced density by harvesting, and changes in primary producers that can have a natural cause (algal blooms) or be indirectly linked to harvesting too, as it was seen that fishing led to higher abundance of primary producers (Frank et al. 2005). Hence, size-selective harvesting provides an ideal context to explore genotype-by-environment interactions on the gut microbiome but have been largely understudied so far.

In fisheries science, laboratory size-selection experiments are commonly used to mimic the evolutionary consequences of size-selective fishing, while controlling for phenotypic plasticity (Conover and Munch 2002; Uusi-Heikkilä et al. 2015). Recently, Renneville et al. (2020) performed a size-selection experiment using medaka (*Oryzias latipes*) as a model species. Native to East Asian countries, medaka is a small cyprinodont fish (adult length = 32 mm) that has a short generation time and is easily reared in the laboratory, making it an ideal species for selection experiments (Ruzzante and Doyle 1993, Renneville et al. 2020, Bouffet-Halle et al. 2021). The species is omnivorous with an animal-based diet preference but can also feed on diatoms and filamentous algae (Edeline et al. 2016). The size-selection procedure consisted of mimicking either fishing mortality where only small-bodied fish were allowed to reproduce (small-breeder SB line), or a more natural mortality regime favoring the reproduction of large-bodied fish (large-breeder LB line) (Reneville et al. 2020, Le Rouzic et al. 2020). As expected from the literature (Stearns 1992, Conover and Munch 2002), the LB and SB lines evolved opposite life-history traits and behaviors: small-breeder medaka grew slower, matured earlier and were less efficient foragers than the large-breeder medaka (Diaz Pauli et al. 2019, Evangelista et al. 2021). The logical next step is to examine to what extent changes in the gut microbiota of LB and SB medaka are driven by the interaction between fisheries-induced changes in both evolution (life-history shift) and environmental conditions (reduced fish density and concomitant increased resource availability).

In the present study, we assessed how the genetic background of the two medaka lines interacted with population density and light intensity (used to modulate primary production) to shape the gut microbiome composition and diversity. For this, SB and LB medaka were introduced into replicated pond mesocosms for three months. Based on life-history and foraging traits divergence between the two lines (Diaz Pauli et al. 2019, Evangelista et al. 2020, Evangelista et al. 2021), and based on the assumption that such traits are key drivers of gut microbiome variation in teleosts (Talwar et al. 2018), we hypothesized divergent gut microbial communities between SB and LB medaka. We further hypothesized that gut microbiome differences between the two lines would be more pronounced under suboptimal conditions, i.e. when access to food resources is limited (Reese and Dunn 2018, Varg et al. 2021). Finally, because microbiome diversity could be important for host fitness, we evaluated whether fitness proxies (i.e. somatic growth rate and body condition) were associated with variation in gut microbiome diversity (Bolnick et al. 2014b).

## Methods

### Size-dependent selection and fish rearing

The two medaka lines were size-selected over 10 generations under identical laboratory conditions. Specifically, medaka were kept in 3-L tank at similar density (14 – 17 fish per tank, 20 tanks per line per generation; Le Rouzic et al. 2020, Renneville et al. 2020), and at the same temperature (26°C) and photoperiod (14 h Light / 10 h Dark). They were fed *ad libitum* with a mixed diet of dry food and living brine shrimp *Artemia salina* and/or *Turbatrix aceti*. These standardized environmental conditions ensured that phenotypic differences among the selected lines reflected a genetically-based, evolutionary divergence in response to size-selective harvesting alone (Le Rouzic et al. 2020, Renneville et al. 2020). In addition, offspring from each breeding pair were transferred to the same tank, so one was able to keep track of individual pedigrees during the whole selection experiment (Renneville et al. 2020).

The selection procedure consisted in removing the largest or the smallest breeders, thus producing two lines with distinct life-history strategies: the small-breeder line where only small-bodied individuals were allowed to reproduce, and the large-breeder lines. At each generation, on average 88% of fish were removed per line. Size selection was both family- and individual-based. At 60 day-post-hatching (dph), among a total of at least 20 families per line (one tank per family to keep track of individual pedigrees), the 10 families with the largest (large-breeder line) or smallest (small-breeder line) average standard body length (SL) were kept. At 75 dph, individuals within each of the selected families were measured and the largest-bodied (large-breeder line) or the smallest-bodied (small-breeder line) mature males (n = 2 or 3) and females (n = 2 or 3) were used as breeders for the next generation (further details available in Renneville et al. 2020). On average, at 75 dph, SL was 20.7 mm in small breeders and 22.0 mm in large breeders (a 5.7 % difference), and the probability of being mature was 91.7% in small breeders and 77 % in large breeders (a 18.0 % difference) (Renneville et al. 2020).

In June 2017, experimental populations were created using fish from the eleventh generation. Specifically, for each line, 180 mature fish (initial standard body length: mean ± SD; SL_i_ in small-breeder = 18.9 mm ± 1.4; SL_i_ in large-breeder = 19.4 mm ± 1.4; ANOVA: *F*_1, 358_ = 13.70, *P* < 0.001) were selected to generate 24 experimental populations composed of individuals from the same line (48 populations in total), but from distinct families to limit inbreeding (mean kinship coefficient = 0.23 ± 0.1 and 0.17 ± 0.1 SE in LB and SB lines, respectively; further details available in Le Rouzic et al. 2020). Selected fish were anaesthetized with MS-222 and marked using visible implant elastomer (VIE; Northwest Marine Technology, Shaw Island, WA, USA) to render each fish individually identifiable and to allow the calculation of growth-related traits. Fish from the same experimental population were pooled in a 3 L tank and maintained at the laboratory until the beginning of the experiment when they were released into an outdoor mesocosm (48 mesocosms in total).

### Outdoor mesocosm experiment

The outdoor experiment was conducted at the CEREEP-Ecotron Ile de France (Saint-Pierre-les-Nemours, France; <u>cereep.bio.ens.psl.eu</u>) using 48 mesocosms (500 L, 0.8 m deep, 1.0 m diameter) arranged in 5 blocks. All mesocosms were filled simultaneously from 4 to 6 April 2017 with a mix of dechlorinated tap water (100 L) and oligotrophic water from a local pond (300 L). The pond water was pre-filtered through 150 μm mesh to remove large benthic invertebrates, zooplankton and debris. The mesocosms were supplied with 2 L of mature sediment mixture including benthic invertebrates (mainly Ephemeroptera and Chironomidae larvae, Planorbidae, Hydrachnidia, Nematoda and Ostracoda) and 2 L of homogenized mixture of zooplankton (Copepoda and Cladocera) collected from local ponds. In each mesocosm, two floating shelters made of wool threads (30 cm length) provided spawning substrate and two floating brushes made of plastic threads provided protection for larvae. Each mesocosm was then covered with a shading net (see details below) and given 3 months to mature before fish were introduced. On 12 June, all mesocosms were enriched with 2 mL of a liquid mixture of 0.32 μg P L^-1^ as KH_2_PO_4_ and 6.32 μg N L^-1^ as NaNO_3_ to favor primary production.

On 4 July 2017, large- and small-breeder fish were released into the outdoor mesocosms under contrasting environmental conditions. The experiment consisted of a 2 × 2 × 2 full factorial design with size-selected line (LB and SB) crossed with density (high HD and low LD) and light intensity (high HL and low LL). Each treatment combination was replicated six times (48 mesocosm in total; Fig. 1). High- and low-density treatments consisted of twelve (3.2 mg fish L−1 ± 0.3 SD) and three (0.9 mg fish L−1 ± 0.1 SD) fish per mesocosm, respectively, with a female biased sex ratio of 2:1. Light intensity was manipulated using shade nets with different mesh size that allowed the passage of 92% (high light intensity HL) and 70% of ambient light (low light intensity LL). Light supply was used to modulate primary production while avoiding too high growth of filamentous algae. This factorial design resulted in a total of 8 treatment combinations (Fig. 1), each with 6 replicates.

**Figure 1.**
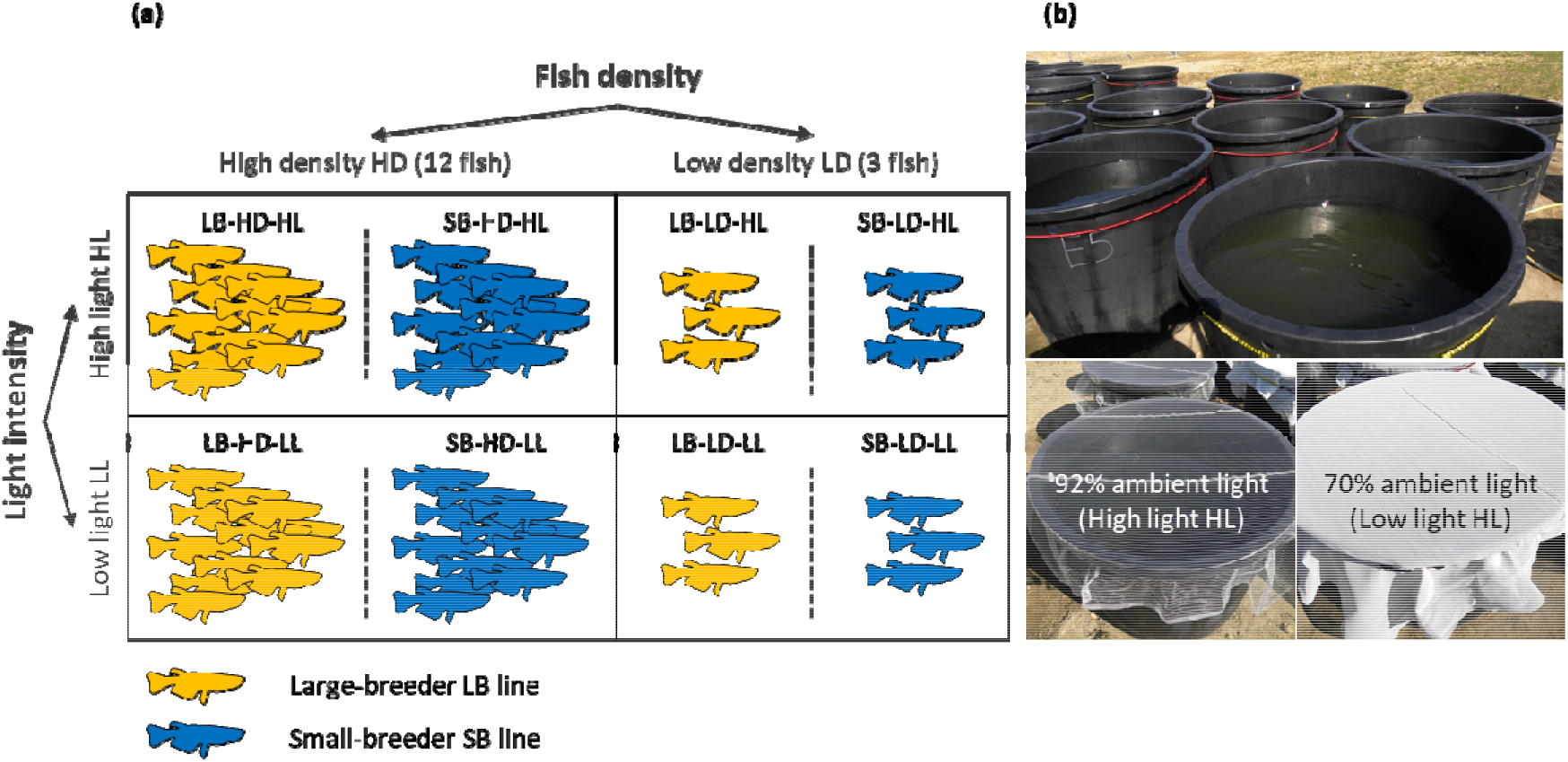
**(a)** Design of the mesocosm experiment used to test the effects of Line × Density and Line × Light intensity on fish gut microbiome. Fish from the Large-breeder (LB) line are in orange and fish from the Small-breeder (SB) line are in blue. **(b)** Pictures of the outdoor mesocosms (upper picture) and shade nets used to manipulate light intensity (lower pictures).

### Gut microbiome sampling and growth-related trait measurements

On 22 September 2017, marked fish were recaptured with hand nets (survival rate = 92%). A total of 126 marked fish were randomly and homogeneously sub-sampled among the mesocosms (number of fish per treatment: mean ± SD = 15.8 ± 0.9, min = 14, max = 17; number of fish per mesocosm: mean ± SD = 2.6 ± 0.6, min = 1, max = 4). After 24 hours fasting, each selected fish was measured for final standard length (SL_f_ ± 1 mm), weighed (W_f_ ± 1 mg), euthanized using MS-222 and dissected using disposable laboratory-grade razor blades. The whole intestine (including potential remaining content because of small size) was sampled and flash-frozen in liquid nitrogen for up to 5 hours, then stored in a -80LJ freezer until DNA extraction. To limit contaminations during dissection, working environment and dissection tools were sterilised between each individual. Scalpels were thoroughly washed using 96% laboratory grade ethanol.

Body condition of each selected individual was calculated using the residuals of the relationship between log_10_W_f_ and log_10_ SL_f_. The somatic growth rate (mm month^-1^) of each selected fish was calculated as follows:

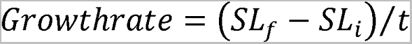

where SL_f_ and SL_i_ are the final and initial standard length, and *t* is the duration of the experiment (3 months).

### DNA extraction and sequencing

After defrosting at room temperature, bacterial DNA from medaka gut samples was extracted using the DNeasy PowerSoil kit (Qiagen, Germany) according to the manufacturer’s instructions. The quantity and quality of purified DNA was checked using a NanoDrop spectrophotometer (Thermo Fisher Scientific, USA). Library preparation for Illumina sequencing was carried out according to the dual indexing protocol described by Fadrosh et al. (2014). This protocol uses the 319F and 806R primer set to amplify the V3-V4 region of the 16S rRNA gene. DNA sequencing was done on an Illumina MiSeq apparatus in 300 bp PE mode. The DNA sequencing was carried out at the Norwegian Sequencing Centre (NSC), and sequence demultiplexing was done using the custom NSC “demultiplexer” software (https://github.com/nsc-norway/triple_index-demultiplexing/tree/master/src), which also removes barcode sequences and heterogeneity spacers. Among the 126 samples, seven displayed amplification failure, and one was removed from the dataset due to mislabelling.

### Bioinformatics analysis

Further sequence data processing was performed using the Divisive Amplicon Denoising Algorithm as implemented in the dada2 v1.16 R-package (Callahan et al. 2016). Taxonomic classification of amplicon sequence variants (ASVs) was carried out using the Ribosomal Database Project v16 training set (Wang et al. 2007). In addition to the fish samples, four DNA extraction and one PCR negative controls (i.e. ultra-pure water instead of DNA) were sequenced and processed as described above. The quality profiles of the negative controls after filtering showed that each control contained very low number of sequences of very poor quality (Evangelista et al., unpublished data). The sequences that did make it through processing (∼290 sequences for all 5 samples combined) corresponded to only 7 unique sequences, each unique to a negative control sample. The taxonomic classification of these 7 unique sequences referred to taxa that are not expected in the fish gut samples (*Moraxella*, *Ralstonia*, *Brevundimonas*, *Massilia*, *Propionibacterium*, *Micrococcus*, and *Knoellia*). This suggests that overall little contamination occurred.

Using the R package phyloseq (v.1.40.0, McMurdie and Holmes 2013), we further filtered the fish gut data in order to remove any contaminant or artefactual sequences. First, ASVs with a Phylum-level assignment probability < 0.80 and those classified as chloroplast DNA were discarded from the dataset. Second, we excluded all ASVs with a total abundance lower than 0.005% of the dataset’s total abundance as they are most likely sequencing errors (Bokulich et al. 2013). Finally, samples with a total sequence reads abundance of < 5000 reads were removed from the dataset (n = 15). The dataset consisted of 103 samples, comprising 627 ASVs for a total of 3,591,039 sequence reads. Sequencing depth ranged from a minimum of 5588 to 85470 reads per sample, with a mean of 34864 reads per sample. Between-sample differences in library sequencing depth were standardized to the median sequencing depth (Appendix S1).

### Statistical analyses

All statistical analyses were run with R v.4.2.1 (R Development Core Team, 2022) using the Family level as taxonomic resolution because it was the best taxonomic level for discriminating (median bootstrap support is 0.52 and 0.89 for Family and Genus level taxonomic assignment, respectively). We used a PERMANOVA to test for differences in community composition according to Line × Density and Line × Light intensity. This was carried out using the *adonis* function in the vegan package (v.2.6.4, Oksanen et al. 2020), by implementing weighted (to take into account information on relative abundance) and unweighted (to take into account taxa presence or absence) UniFrac dissimilarities based on 999 permutations. Statistical tests indicated that there was no deviation from multivariate dispersion (*betadisper* function from vegan), except for the Line × Light intensity factor in the unweighted UniFrac distance PERMANOVA. Nonetheless, sample size was similar between treatment combinations and results should be robust (Anderson and Walsh 2013). Non-metric multidimensional scaling (NMDS) plots with both weighted and unweighted UniFrac distances were used to visualize gut microbiome composition.

Based on these community composition analyses, we agglomerated the data to family level and visualized the relative abundance of normalized data according to the Line treatment. Significant effects of Line or Line × Environment on gut microbiome community composition were further investigated using differential abundance analysis based on the linear discriminant analysis (LDA) on effect size (LEfSe) method. LEfSe was implemented in the microeco package (v.0.12.1, Liu et al. 2021) using a non-parametric Kruskal-Wallis test to detect differences in Family abundance (bootstrap test number = 100, significance threshold = 0.01).

Gut microbiome diversity was estimated using the first three Hill numbers (*^q^* D; Chiu and Chao 2014, Alberdi and Gilbert 2019a) calculated using the R package hilldiv (v.1.5.1, Alberdi and Gilbert 2019b): *q* = 0 (species richness), *q* = 1 (the exponential of Shannon’s entropy index) and *q* = 2 (the inverse of the Simpson’s diversity index). Linear models were used to test the effect of Line × Density and Line × Light intensity on each Hill number.

When significant, the interactions were further investigated using post hoc Tukey’s pairwise comparison using the emmeans package (v.1.8.1.1, Lenth 2021). Spearman correlations (adjusted for multiple testing using false discovery rate (fdr) correction) were used to test for associations between bacterial richness and diversity (i.e. the first three Hill numbers) with medaka growth-related traits (i.e. body condition and somatic growth rate), as well as with medaka final standard length. These correlations were performed using the *corr.test* function from the psych package (v.2.2.9, Revelle 2021). Finally, we applied a Canonical Correlation Analysis (CCA) to explore the relationships between gut microbiome composition and medaka’s traits. CCA was conducted using the mixOmics package (v.6.23.4, Rohart et al. 2017). Results from the CCA were visualised using a correlation circle plot.

### Results Composition of the whole gut microbiome

After standardization of the data, we identified a total of 3,189,868 sequence reads (mean = 30,969 reads per sample) and 627 ASVs for 103 samples. As expected, dominant phylum of the medaka gut microbiome included Proteobacteria (60.5% and 59.2% of reads in LB and SB medaka, respectively), followed by Bacteroidetes (2.5 and 2.4%, respectively), Verrucomicrobia (2.0 and 3.2%, respectively) and Actinobacteria (2.2 and 3.9%, respectively). Bacterial communities were also characterized by large relative abundance of Cyanobacteria (29.2 and 28.8%, respectively), while Firmicutes represented only 0.7% and 0.5% of reads in LB and SB medaka, respectively (Fig. S2).

### Genotype-driven variation in gut microbiome

The two medaka lines had distinct gut microbial communities (weighted UniFrac distance PERMANOVA: *F* = 2.30, *P* = 0.025, R^2^ = 0.022; Fig. 2a). Overall, 5 families (i.e. *Aeromonadaceae*, *Neisseriaceae, Family_II*, *Rhodobacteraceae* and unspecified *Cyanobacteria*) dominated the gut microbiome of all samples, comprising together 62% and 59% of the total bacterial relative abundance of LB and SB medaka lines, respectively (Fig. 2b). Of these families, the relative abundance of both *Aeromonadaceae* and *Neisseriaceae* were significantly higher in LB than in SB medaka (LEfSe; *P* = 0.001 and *P* = 0.016, respectively; Fig. 2c), while *Rhodobacteraceae* was more abundant (relative abundance) in the gut of SB than LB medaka (LEfSe: *P* = 0.017; Fig. 2c). The two other families did not significantly differ between the two lines. Unweighted UniFrac distance PERMANOVA revealed no significant difference between SB and LB medaka (*F* = 1.32, *P* = 0.109, R^2^ = 0.013), indicating that low-abundance taxa did not differ between the two lines.

**Figure 2.**
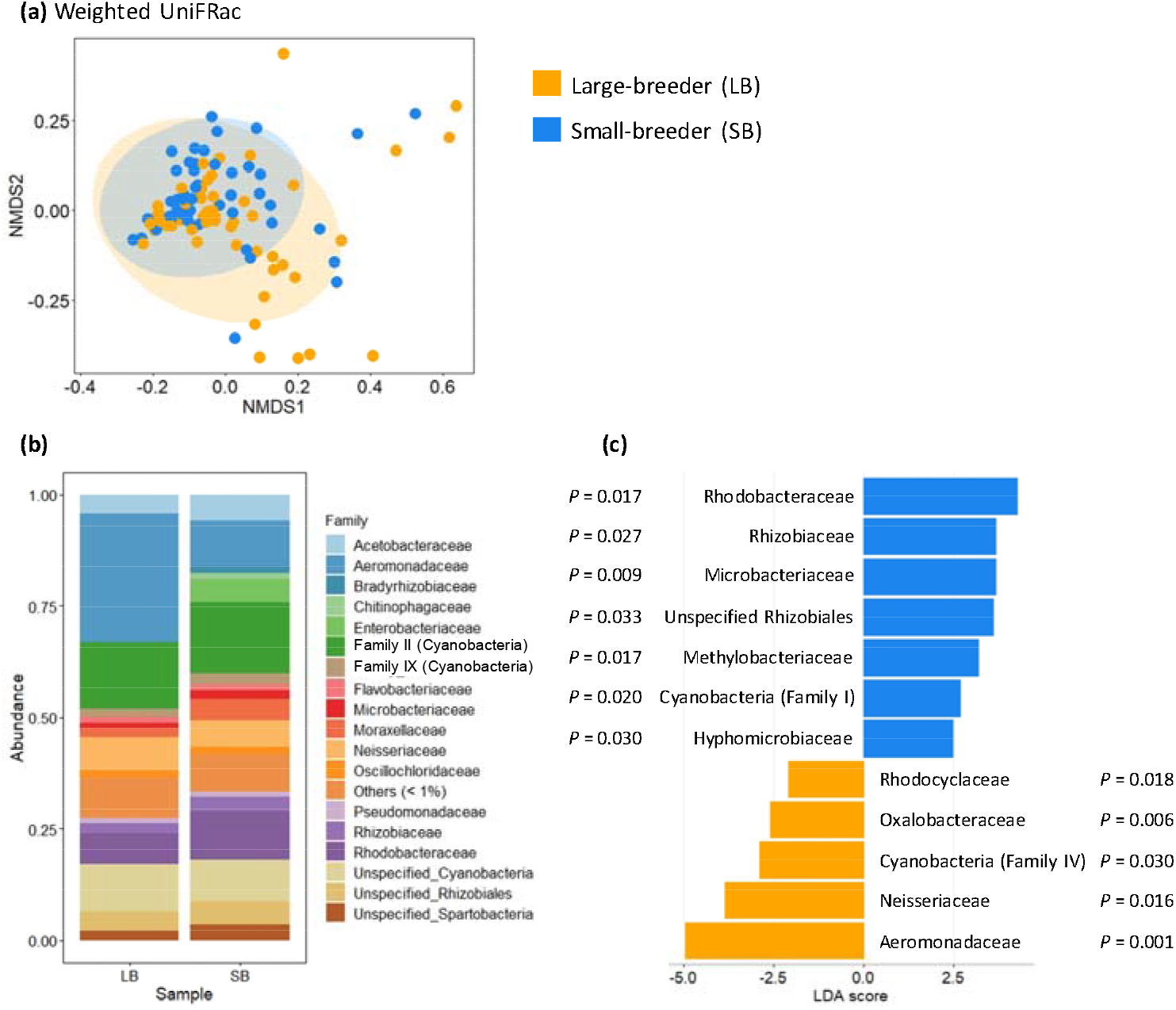
**(a)** NMDS ordination (NMDS stress = 0.13) of variation in bacterial community composition of fish from the Large-breeder (LB; orange dots) and Small-breeder (SB; blue dots) lines. Data represent ordination based on weighted UniFrac distances among the 103 fish individuals. **(b)** Gut microbial taxonomic composition (Family level) of fish according to its selective background (Large-breeder line, LB; Small-breeder line SB). **(c)** Linear discriminant analysis (LDA; LDA score > 2) effect sizes representing the twelve ASVs (Family level) that significantly differ in relative abundance between the Large-breeder (orange) and Small-breeder (blue) lines.

Differential relative abundance analyses showed that *Family IV*, *Oxalobacteraceae*, and *Rhodocyclaceae* were significantly more abundant in the gut of LB than SB medaka (*P* = 0.030, *P* = 0.006, *P* = 0.018, respectively). In contrast, the gut microbiome of SB medaka had more *Microbacteriaceae* (*P* = 0.009), *Rhizobiaceae* (*P* = 0.027), *Rhizobiales* (*unspecified Family*, *P* = 0.033), *Methylobacteriaceae* (*P* = 0.017), *Family I* (*P* = 0.020) and *Hyphomicrobiaceae* (*P* = 0.030) than that of LB ones (Fig. 2c).

### Environment-driven variation in gut microbiome

Weighted and unweighted UniFrac distance PERMANOVA showed no effect of light intensity (F = 1.23, *P* = 0.265, R^2^ = 0.012; F = 1.45, *P* = 0.061, R^2^ = 0.014) and fish density (F = 1.35, *P* = 0.195, R^2^ = 0.013; F = 1.31, *P* = 0.111, R^2^ = 0.013) on the gut microbial community (Fig. 3). Similarly, the gut microbiome diversity of medaka (species richness, exponential Shannon or inverse Simpson) were not influenced by the environmental conditions alone (Table 1).

**Figure 3.**
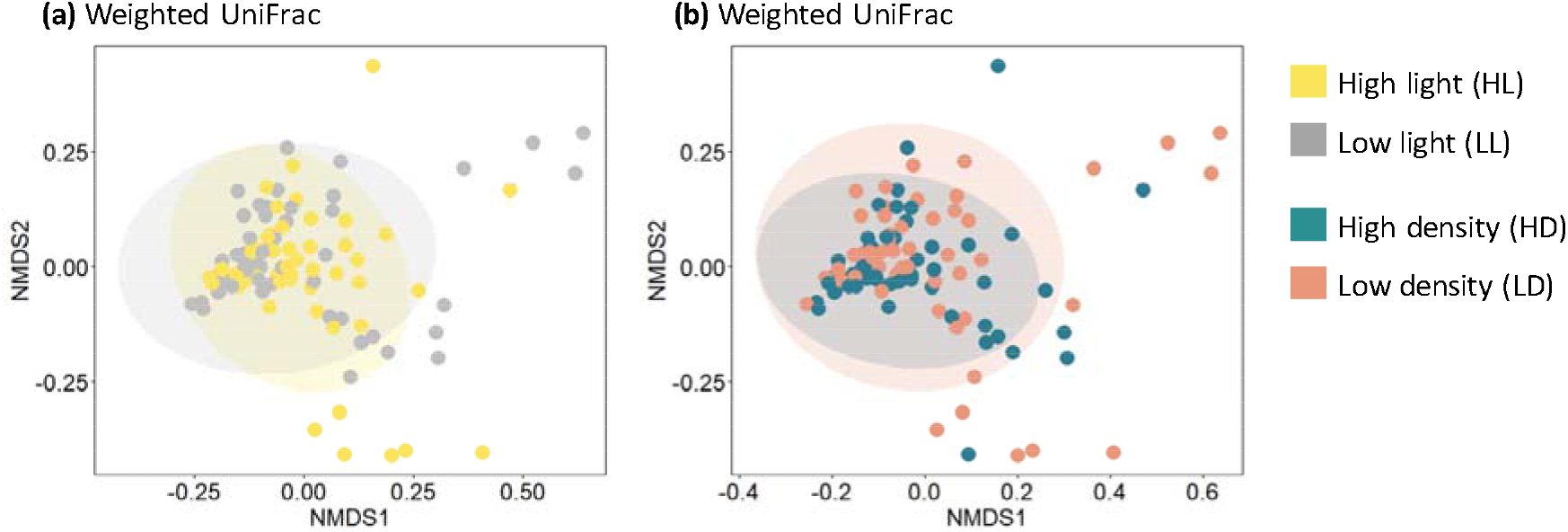
NMDS ordination (NMDS stress = 0.13) of variation in bacterial community composition **(a)** in low or high light intensity (grey and yellow dots, respectively), **(c)** in low or high fish density (pink and green dots, respectively. Data represent ordination based on weighted UniFrac distances among the 103 fish individuals.

**Table 1.**
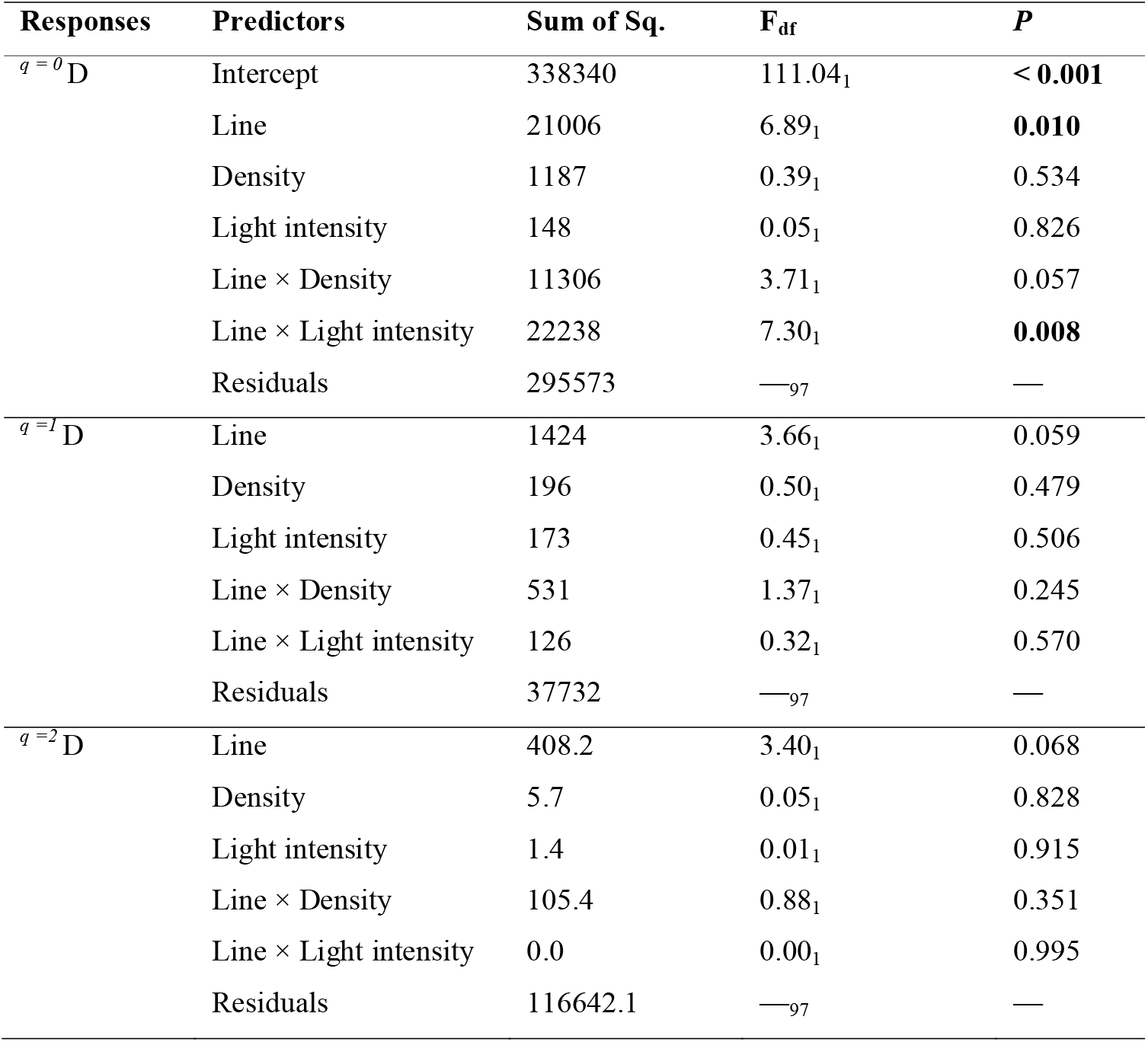
Analysis-of-variance table derived from the linear models used to assess the effect of size-selected line, fish density, and light intensity on gut microbiome diversity (*^q= 0^* D, *^q=1^* D and *^q=2^* D). Significant P values are highlighted in bold.

### Gene-by-environment interaction on gut microbiome

Weighted UniFrac analyses showed that the interactions between medaka line and the environment (i.e. Line × Density and Line × Light intensity) had no significant effect on gut microbiome composition (PERMANOVA: *F* = 1.08, *P* = 0.330, R^2^ = 0.010 and *F* = 0.82, *P* = 0.561, R^2^ = 0.001, respectively). In contrast, unweighted UniFRac analyses revealed that Line × Light intensity significantly influenced the gut microbial composition (*F* = 1.66, *P* = 0.020, R^2^ = 0.016). Indeed, the gut microbiome of LB medaka depicted greater compositional diversity than that of SB medaka in the high-light treatment, while the opposite pattern was observed in the low-light treatment (Fig. 4a).

**Figure 4.**
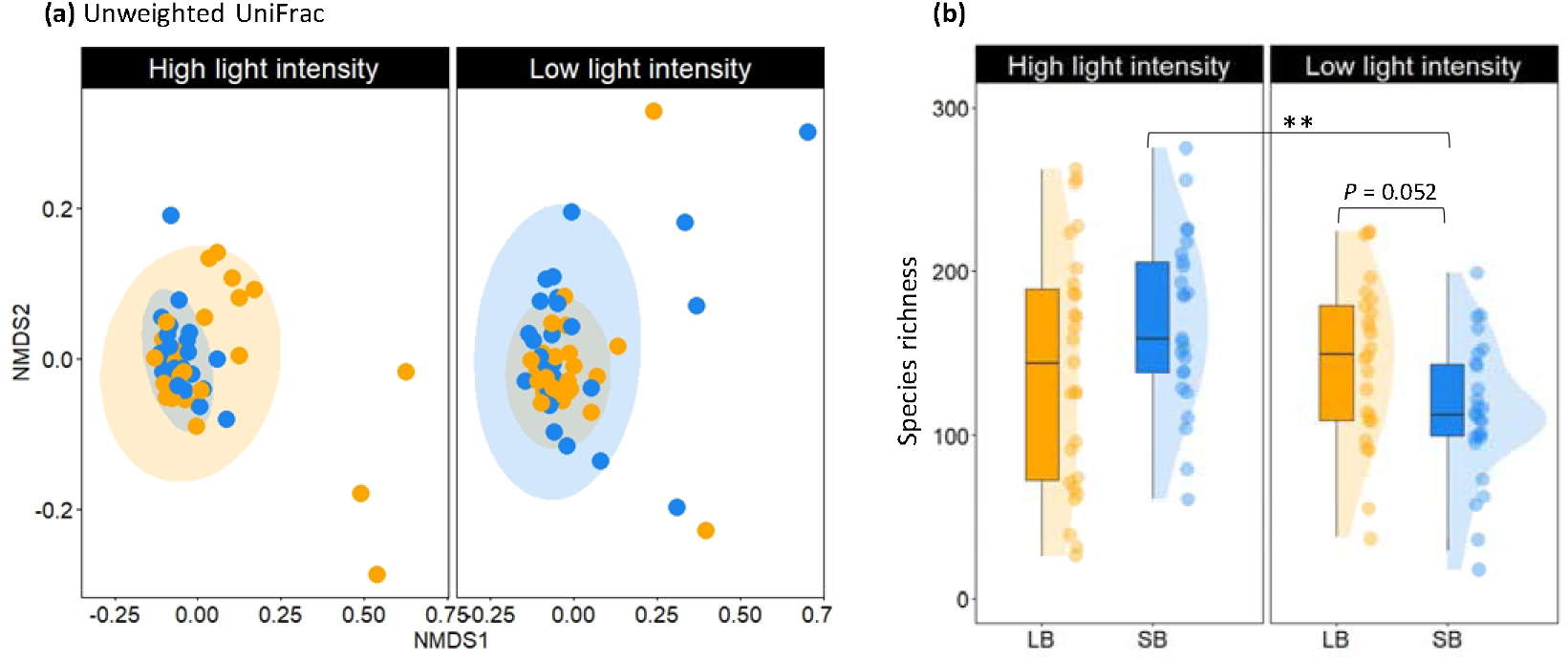
**(a)** NMDS ordination (NMDS stress = 0.15) of variation in bacterial community composition of SB (blue) and LB (orange) fish in the different light conditions. Data represent ordination based on unweighted (d;) UniFrac distances among the 103 fish individuals. **(b)** Raincloud plot showing the light-intensity effect of Line on bacterial richness (*^q= 0^* D). Dots represent the fish (n = 103), boxplots and half violin plots illustrate the probability density of the data.

Gut microbial richness was modulated by the interaction between the medaka line and light intensity (Table 1). Specifically, SB medaka had a higher bacterial richness (when q = 0) in the high-light intensity compared to the low-light treatment (Line × Light: Tukey post hoc: *^q= 0^* D_SB-HL_ vs. *^q= 0^* D_SB-LL_: t_97_ = 3.59, *P* < 0.001; mean *^q= 0^* D ± SE: 169 ± 11 and 114 ± 8, respectively), while bacterial richness of LB medaka did not change with light variation (Fig. 4b; *^q= 0^* D_LB-HL_ = 141 ± 14, *^q= 0^* D_LB-LL_ = 145 ± 10). However, LB medaka seemed to have a higher bacterial species richness than SB medaka, but only in the low-light treatment (Line × Light: Tukey post hoc: *^q= 0^* D_LB-LL_ vs. *^q= 0^* D_SB-LL_: t_97_ = 1.97, P = 0.052). Other metrics of gut microbiome diversity (when *q* = 1 or 2) were not influenced by the interaction between Line and environmental conditions (Table 1).

### Lack of correlation between microbiome and growth-related traits

No significant correlation was observed between gut microbiome diversity (estimated using the first three Hill numbers) and medaka’s body condition, somatic growth, and final standard length (Spearman correlations: adjusted *P* > 0.876 for all; Appendix 3). Correlations between medaka’s traits and gut microbial Families were not detected (Fig. 5).

**Figure 5.**
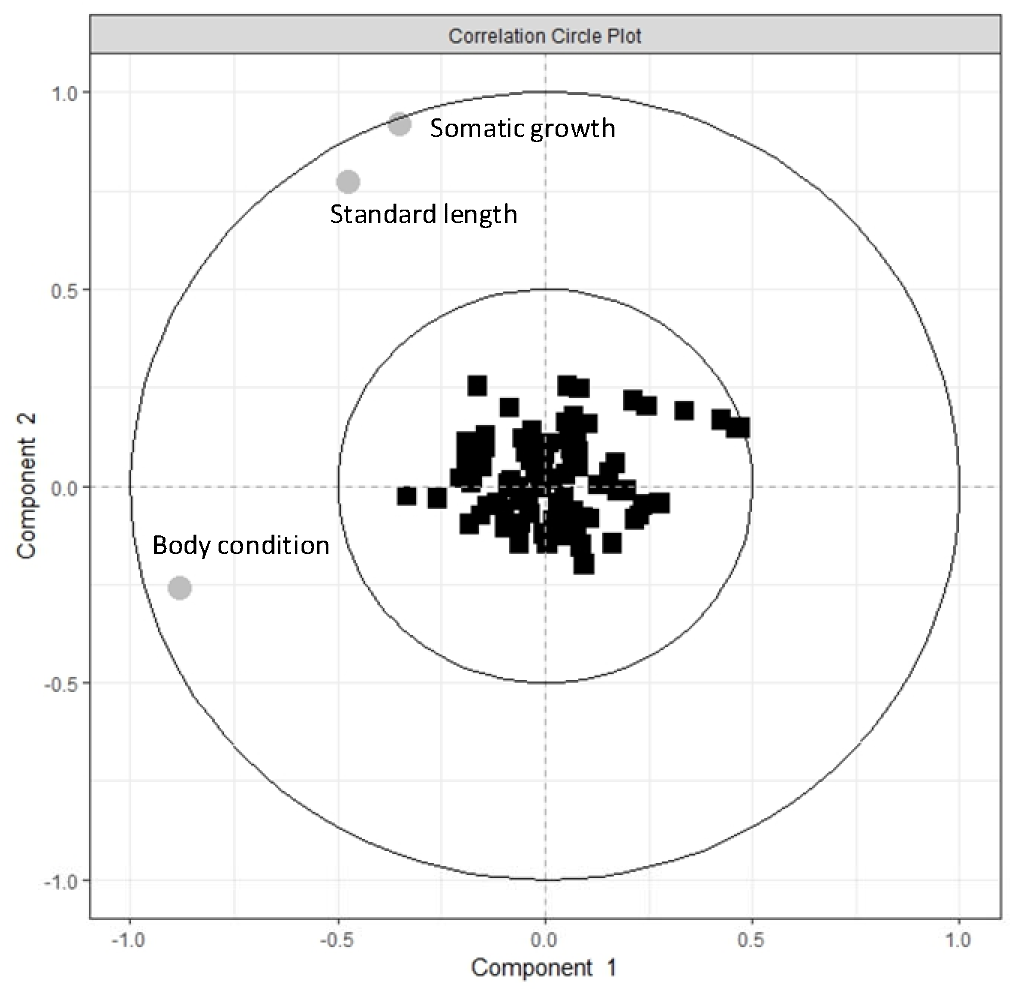
Correlation circle plot from the CCA applied between medaka’s traits (grey circles) and gut microbial composition (Family level, black squares). The farther from the center a bacterial family or traits is, the greater the association with the component. Variables projected in the same direction of the plot are positively correlated.

## Discussion

Due to its pivotal role for host fitness and health, there has been a growing research interest in the factors driving gut microbiome variation in animal species. Yet, studies focusing on the genotype-by-environment effects on the gut microbiome of fish remain limited (Sevellec et al. 2014, Piazzon et al. 2020). Using size-selected medaka lines (large-breeder LB and small-breeder SB) in a pond mesocosm experiment, we found that the composition of the core gut microbial community differed between the two lines. In addition, differences in rare taxa (unweighted UniFrac) between the lines were modulated by light intensity, with more dispersion for LB medaka than SB medaka in the high-light treatment and the opposite pattern in the low-light treatment. The microbiome richness of SB medaka was influenced by light intensity, while that of LB remained unchanged regardless of the environmental conditions. Together, this is consistent with our prediction that evolutionary changes due to size-selective harvesting have the potential to shape the gut microbiome assemblage within harvested populations. Our results also suggest that the interaction between the genetic background of medaka (i.e. the selected line) and the environmental conditions could be important, as previously reported in the gut microbiome communities of two guppy ecotypes, sampled across different streams (Sullam et al. 2015). However, contrary to our prediction, variation in microbiome diversity or composition was not associated with any of the measured growth-related traits of adult medaka.

Our findings confirm observations from literature showing that genotype could drive gut microbiome composition among fish groups (Sevellec et al. 2018, Small et al. 2019, Smith et al. 2015). In our case, the largest difference in relative read abundance was found for the *Aeromonadaceae* family which was almost 3 times more abundant in LB than in SB medaka (28.9% and 10.1%, respectively). Our previous findings from the same pond experiment suggested that adult and juvenile LB medaka foraged more on benthic prey hidden in the sediments than the SB medaka (Evangelista et al. 2021). Although further investigation is required to back-up this hypothesis, increased relative abundance of *Aeromonadaceae* in LB might reflect distinct foraging strategies between the two lines as *Aeromonadaceae* are known to be facultative aerobes and are mainly found in anoxic sediments (Tomás 2012, Laviad and Halpern 2016). Additionally, some *Aeromonas* display cellulolytic activity, which can be useful for the digestion of plant-based diet (Li et al. 2016). One could hypothesize that a higher proportion of *Aeromonadaceae* in LB medaka could be associated with a more omnivorous feeding habit compared to SB medaka (Liu et al. 2016). In addition, *Neisseriaceae* found in greater quantities in LB was also found to be more abundant in rainbow trout fed with fish-based meal as to opposed to those fed with insect meal (Terova et al. 2021), potentially suggesting that LB fish predate more on fish larvae than SB medaka. Interestingly, *Rhodobacteraceae*, which were found in higher proportion in SB medaka here, has been associated with higher levels of environmentally induced stress in the host (Pootakham et al. 2019 and references herein). This might add another line of evidence about the lesser capacity of SB medaka to cope with a less favourable surrounding environment, as previously reported (Evangelista et al. 2020, Evangelista et al. 2021). Altogether, our results suggest that even if the gut microbiome composition between the two lines differs, the mere description of microbiome diversity and composition is not sufficient to precisely identify the underlying causes of gut microbiome variation between SB and SL medaka. In fact, more targeted diet manipulation experiments between lines (e.g. Sullam et al. 2015) would be required in order to clearly identify whether adaptation to size selection could directly affect the gut microbiome, or indirectly through changes in diet.

Under low-light intensity, the gut microbiome of SB medaka showed a 34% decrease of bacterial richness compared to the high-light intensity treatment, and also presented a somewhat lower richness than in LB medaka, though this difference was not significant (P = 0.052). This suggests that the gut microbiome diversity of fish selected for earlier maturation and slower growth rate (as is often the case with size-selective harvesting) can be reduced under sub-optimal environmental conditions. The underlying mechanisms of such genotype-by-environment effects are hard to pinpoint and can be the results of many interacting factors – light-induced changes in the SB medaka behaviour or physiology, combined with changes in diet or in the bacterial composition of water, ultimately altering the gut microbiome. However, as we did not sequence the microbiome from the water used in the experimental ponds, we are not able to assess the extent to which variation in water composition or even potential co-amplification of bacterial taxa from the environment (Talwar et al. 2018) could influence our results. In term of bias, previous studies suggest that fish gut microbiome composition is not influenced by the bacterial composition of the surrounding waters (Schmidt et al. 2015, Wang et al. 2018), thus implying water should not be a confounding factor in the context of our study. The present study focused on the deterministic processes that influenced the gut microbiome composition of medaka, but it is important to note that stochastic processes may also explained substantial variability in the bacterial composition of medaka (Jones et al. 2022).

Whether high microbial diversity matters for the host remains a central question in microbiome studies. For instance, Bolnick et al. (2014b) found a positive effect of the gut microbial diversity on the body condition of laboratory-reared stickleback (*Gasterosteus aculeatus*), but no such effect on condition of wild stickleback. In our experiment, gut microbiome diversity was not associated with growth-related traits of adult medaka, perhaps because low diversity does not entail the loss of essential microbiome-mediated functions. Therefore, the lack of associations could simply reflect our yet limited understanding about the taxonomic identity and functional role of gut bacteria in non-model organisms. Thus, changes in bacterial diversity might be associated with either positive or negative impact for the host, according to the degree of decoupling between taxonomic identity, functional role and the environmental context (Bolnick et al. 2014b). Overall, our results highlight the fact that our perception about gut microbiome benefits is also probably biased by data based on a very limited range of species (Hammer et al. 2019). Nonetheless, variation in the microbiome can impact the digestive capacity and body condition, and this diversity may act as an underlying mechanism for phenotypic plasticity of the host (Alberdi et al. 2016). How changes in bacterial diversity translate into functional changes will require further investigation.

Our study reveals that the gut microbiome of fish can be influenced by interactions between their genetic background and the environment. Studying genotype-by-environment interactions on the gut microbiome may bring new perspectives into the role of microbiomes in eco-evolutionary dynamics, as changes in gut microbial communities could potentially reverberate into changes in ecosystem functioning and services (Graham et al. 2016, Dutton et al. 2021), including fisheries productivity (Gallo et al. 2020, Diwan et al. 2021). As the demand for fish for human consumption is increasing, we also claim that more research is needed to enhance our understanding of the possible effects of fisheries-induced evolution on the gut microbiome. Comparison of the gut microbiome of fish in relation to different management strategies (e.g. balanced versus size-selective fishing versus protected areas) may reveal important mechanisms influencing populations’ adaptability and resilience, and thus help restoring highly impacted fish stocks (Gallo et al. 2020). It is also important for future studies to reveal the functional consequence of changes in gut microbiome (Ghanbari et al. 2015, Tarnecki et al. 2017), especially functions directly responsible for fish behaviour and fitness so that the target preservation of highly beneficial gut microbiomes within harvested populations could be incorporated as part of more sustainable fisheries practices.

## Acknowledgments

We are grateful to Clémentine Renneville and Arnaud Le Rouzic for initiating the medaka lines; David Carmignac and Romain Péronnet for their help maintaining the fish; Julia Dupeu, Anders Herland and Jacques Meriguet for field assistance; the platform PLANAQUA and the CEREEP Ecotron Ile-De-France for the access to the experimental facilities; and Natacha Nikolic for valuable comments on a previous version of this manuscript. We also thank Konstantinos Kormas, Marco Basili, Laetitia Wilkins and one anonymous reviewer for their insightful comments that improved the manuscript.

## Data, script, code, and supplementary information availability

Data and R codes that support the findings of this study are hosted in the Figshare repository (https://figshare.com/s/d5235f25f3d7b15a0e47). SRA accession for sequences data is PRJNA929943

## Ethics

The experiment was approved by the Darwin Ethical committee (case file #Ce5/2010/041) from the French Ministry of Education, Higher Education and Research.

## Author’s contributions

CE conceived and coordinated the study with input from SK, EM and PT; CE, JD and BDP collected the samples; SK and EM carried out the molecular analyses; PT carried out the bioinformatic analyses; CE analyzed the data and wrote the initial draft of the manuscript with input from SK, PT and EM; All authors contributed to revisions and approved the final version of the manuscript.

## Appendix 1: Rarefaction curves

**Figure S1.**
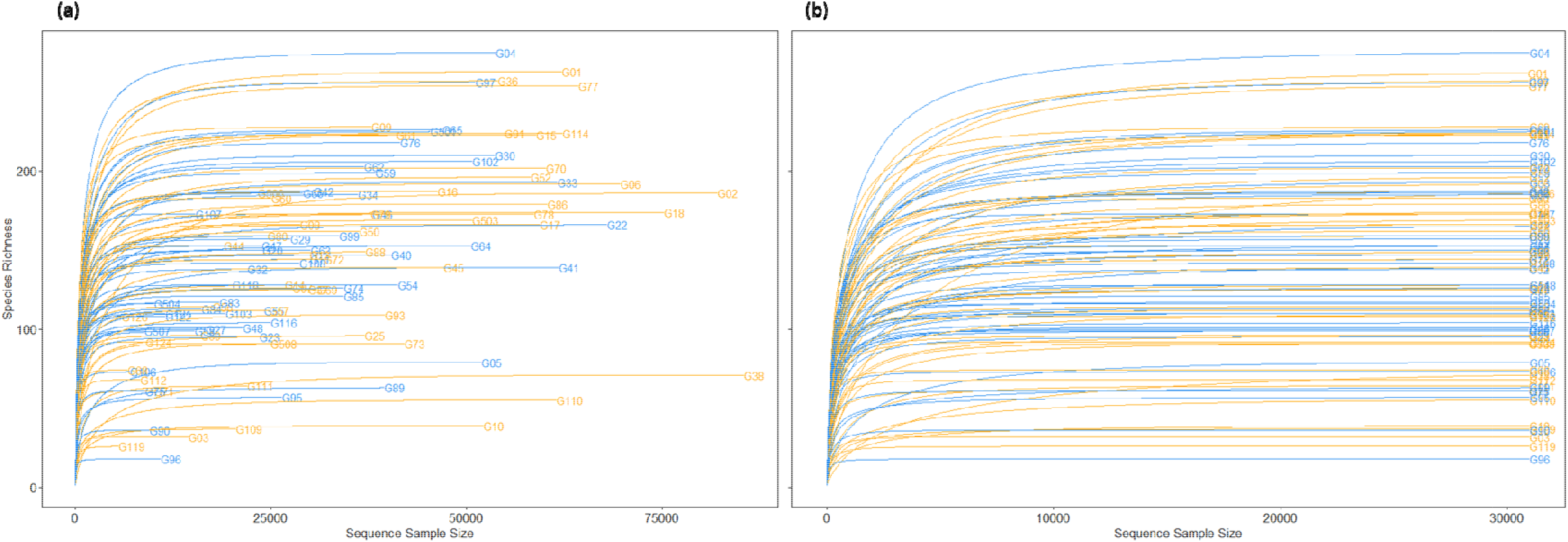
Rarefaction curves showing the number of ASV with increasing number of reads (Sequence sample size) **(a)** before and **(b)** after standardization to the median sequencing depth. Fish from the Large-breeder (LB) line are in orange (n = 52) and fish from the Small-breeder (SB) line are in blue (n = 51).

## Appendix 2: Relative abundance of bacterial phyla in gut samples

**Figure S2.**
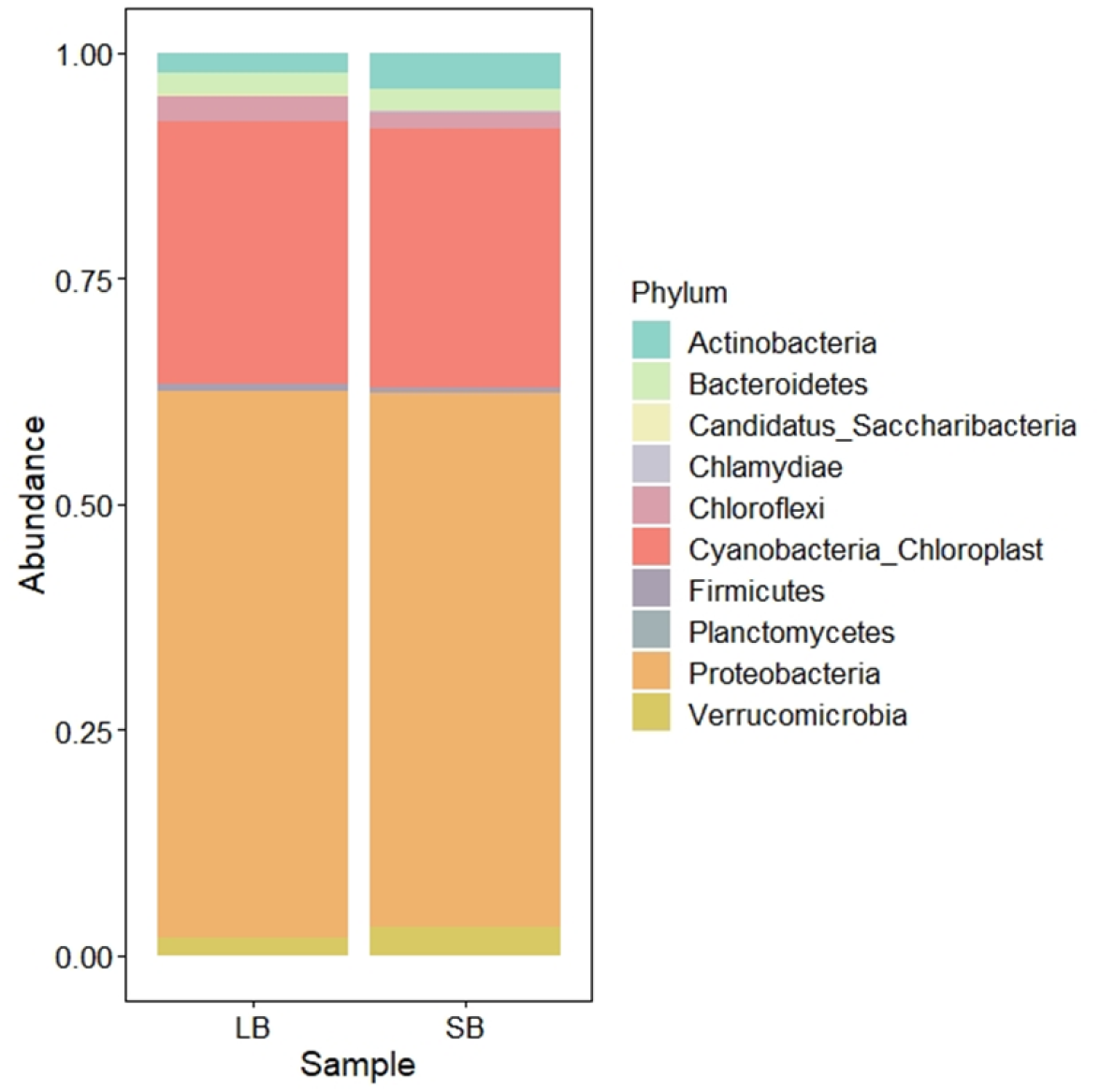
Gut microbial taxonomic composition (Phylum level) of fish according to its selective background (Large-breeder line, LB; Small-breeder line SB).

## Appendix 3: Correlations between medaka growth-related traits and gut microbiome diversity

**Figure S3.**
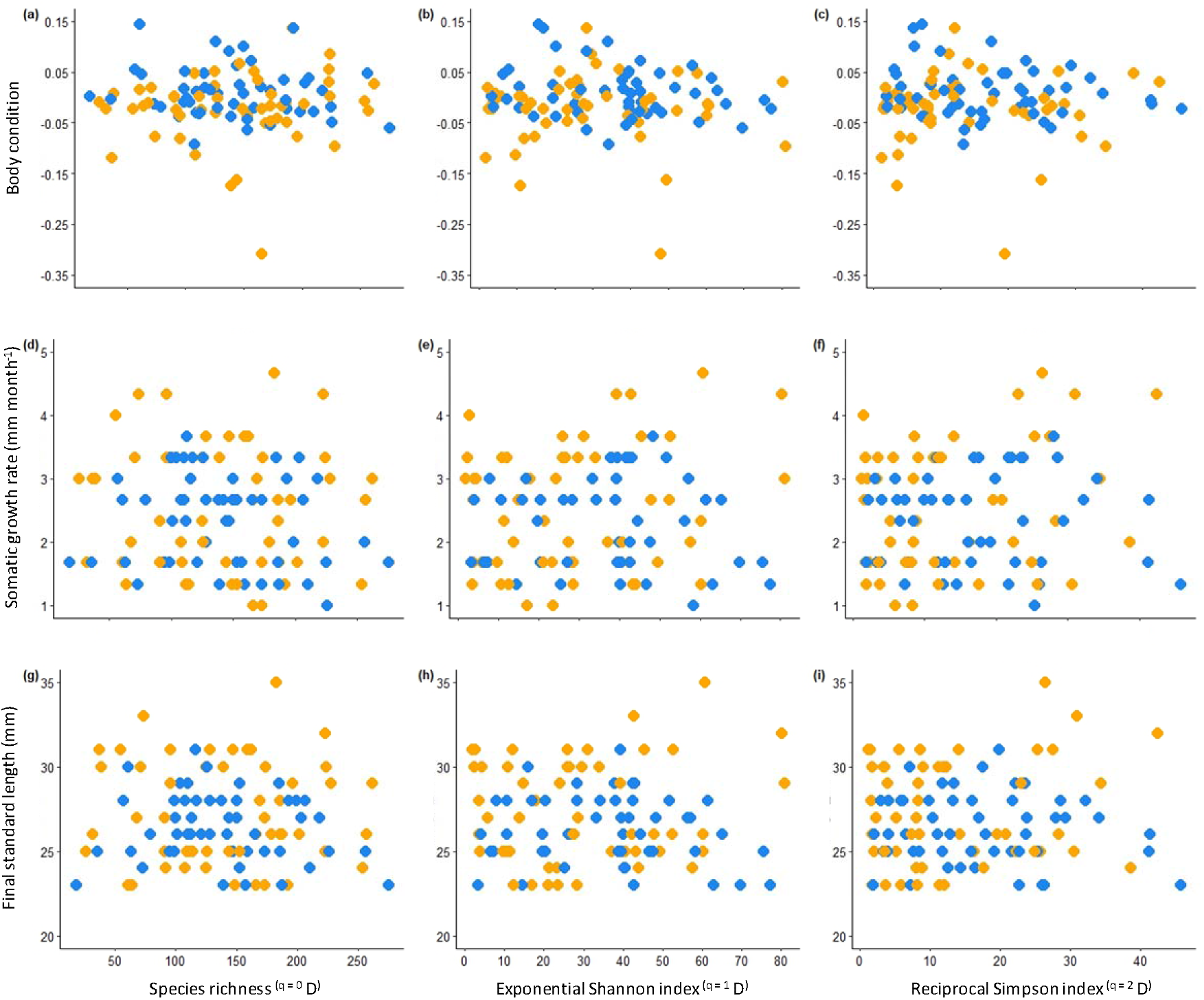
Correlations between gut microbial diversity estimated using the first three Hill numbers (*^q^* D; species richness, exponential Shannon index and Reciprocal Simpson index) and **(a-c)** body condition estimated as the residuals of the relationship between log_10_W_f_ and log_10_ SL_f_, **(d-f)** somatic growth rate [mm month^-1^]) and (**g-i**) final standard length (mm) of fish from the Large-breeder (LB; orange dots, n = 52) and Small-breeder (SB; blue dots, n = 51) lines. None of the correlations were significant.

